# Proteomic changes in bacteria caused by exposure to environmental conditions can be detected by Matrix-Assisted Laser Desorption/Ionization – Time of Flight (MALDI-ToF) Mass Spectrometry

**DOI:** 10.1101/2020.01.24.918938

**Authors:** Denise Chac, Melissa Kordahi, Leandra Brettner, Arushi Verma, Paul McCleary, Kelly Crebs, Cara Yee, R. William DePaolo

**Affiliations:** Department of Pathology, University of Washington, Seattle, WA 98195; Department of Bioengineering, University of Washington, Seattle WA 98195; Division of Endocrinology, Seattle Children’s Hospital I University of Washington, Seattle WA 98115; Department of Medicine, University of Washington, Seattle WA 98195; Center for Microbiome Sciences & Therapeutics, University of Washington, Seattle WA 98195

**Keywords:** MALDI-TOF MS, proteotyping

## Abstract

In the past decade, matrix-assisted laser desorption/ionization time-of-flight (MALDI-ToF) mass spectrometry (MS) has become a timely and cost-effective alternative to bacterial identification. The MALDI-ToF MS technique analyzes the total protein of culturable microorganisms at the species level and produces a mass spectra based on peptides which is compared to a database of identified profiles. Consequently, unique signatures of each microorganism are produced allowing identification at the species and, more importantly, strain level. Our present study proposes that the MALDI-ToF MS can be further used to screen functional and metabolic differences. While other studies applied the MALDI-ToF technique to identify subgroups within species, we investigated how various environmental factors could alter the unique bacterial signatures. We found that genetic and phenotypic differences between microorganisms belonging to the same species can be reflected in peptide-mass fingerprints generated by MALDI-ToF MS. These results suggest that MALDI-ToF MS can screen intra-species phenotypic differences of several microorganisms.

## INTRODUCTION

The MALDI-ToF (matrix-assisted desorption/ionization time-of-flight) mass spectrometry (MS) technology offers a time- and cost-effective method of identifying microorganisms. Compared to previous time-consuming and expensive methods to identify microorganisms based on 16s rRNA or whole genome sequencing, the MALDI-ToF MS provides rapid, accurate and inexpensive identification within minutes via proteotyping^1^. While the MALDI-ToF is limited to culturable microorganisms and public databases, it has been quickly incorporated in the clinical setting and used for diagnosis. Studies have shown that MALDI-ToF MS can equally or even better identify sources of systemic infections^2, 3^, urinary tract infections^4–6^, respiratory tract infections^7^ and intestinal infections^8, 9^. The MALDI-ToF MS technology allows for identification down to the strain level and has been shown to discriminate between strains of methicillin-resistant *Staphylococcus aureus*^10–13^, shiga-toxigenic *Escherichia coli*^14^, clinically relevant strains of Aspergillus species ^15^and many others^1^.

Although many existing methods allow rapid identification of microorganisms to the species level, identification to the more specific “strain” taxon tends to be more challenging, as strains within a single microbial species are often very genotypically and phenotypically similar, despite having different functions. Higher resolution approaches such as molecular genetics are thus commonly employed to identify and characterize strains within a microbial species ^16^. These include pulsed field gel electrophoresis (PFGE)^17^, multilocus sequence typing^18^ (MLST), repetitive extragenic palindromic PCR^19^ (rep-PCR), housekeeping gene (e.g., PheS) sequence analysis^20^, and whole genome sequencing^21, 22^. Each of these approaches has been shown to have adequately high resolution to distinguish microbial strains from one another; however, these approaches are labor- and time-intensive as well as costly techniques that might lack the required rapid high-throughput nature of MALDI-ToF. A big advantage of the MALDI-ToF MS strain typing application would be epidemiologic investigations that require rapid identification of a single strain within a single species to determine the origin and spread of an outbreak in order to mitigate risks to public safety posed by microbial food and water contamination or potential acts of bioterrorism ^16, 23^. Moreover, in an era where microbiome-based diagnostics and therapeutic targets are envisioned^24^, this technology could provide a framework to link disease phenotypes to peptides translated by microorganisms isolated from various biological samples.

The MALDI-ToF MS is an ionization process that became commercially available in the early 1990s. The ionization process involves the mixture of an analyte with a “matrix” solution and the co-crystallization of both matrix and analyte (**Diagram 1 A-B**). During ionization, a laser of a determined UV nm wavelength is fired at the mixture and causes its desorption into a gaseous phase. MALDI ion sources are combined with time of flight (ToF) tubes for ion separation^16^. A peptide mass fingerprint (PMF) is generated per sample and compared to a database of known bacterial species PMFs (**Diagram 1C**). MALDI-ToF looks at whole cell differences as opposed to a method that target a handful of molecules such as qPCR, protein blots, etc. which allows identification of species and strain differences that other molecular techniques would miss. While the technique lacks discriminatory power with some species such as *Enterococucs faecium* and *Staphylococcus aureus*^25^, the MALDI-ToF MS can robustly identify bacterial species in various culturing conditions^26, 27^.

**Diagram 1.**
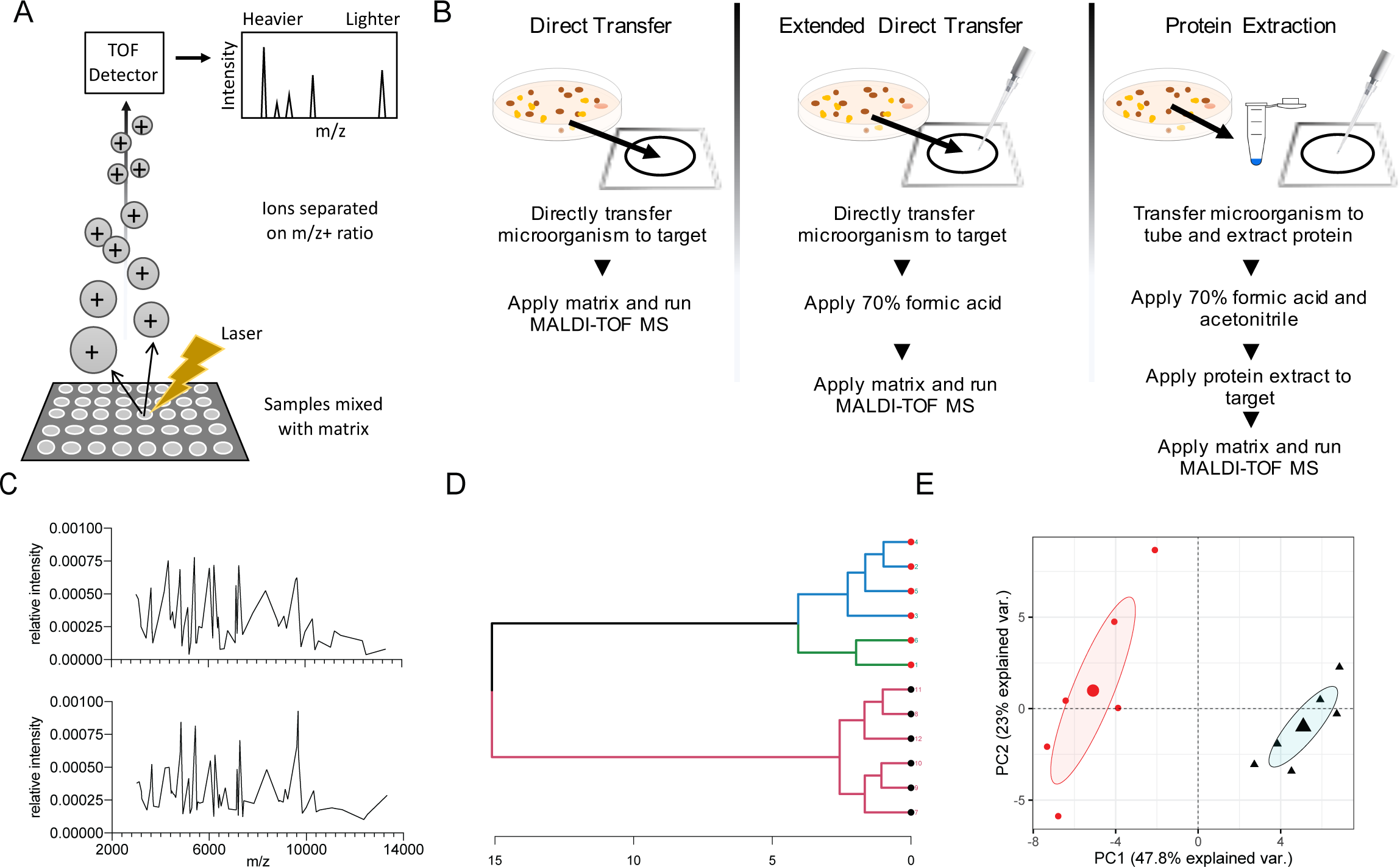
MALDI-ToF MS methodology. A) Schematic of MALDI-ToF MS technology and B) methods of processing as described by manufacturer Bruker. Analysis of peptide-mass fingerprints (PMF) by C) spectra, D) dendrogram and E) principle component analysis (PCA).

In contrast to studies that evaluate the ability of MALDI-ToF MS ability to identify bacterial species regardless of environmental conditions, we propose that the variation of MALDI-ToF MS-produced peaks due to culturing can detect environmental imprints. By comparing PMF peaks and observing the differences in dendrogram and principal component of analysis (PCA) format (**Diagram D and E**), we report that the MALDI-ToF can reliably differentiate between various conditions including oxygen presence, nutrient availability and temperature stress. The MALDI-ToF MS is also able to detect previous environmental conditions after culturing out of the initial environment. Our study shows that the MALDI-ToF can be applied to screening bacterial isolates from various environmental conditions and biological samples in order to see potential functional differences within the same microorganism.

## MATERIALS AND METHODS

### Bacterial strains and growth conditions

DH5α *Escherichia coli* and enteropathogenic *E. coli* were grown on tryptic soy agar (TSA) plates and incubated at 37°C for 24 hours. *Yersinia enterocolitica* 8081 and *Y. enterocolitica INV, YadA, and YscA* knock-out strains were grown on TSA and incubated at room temperature for 48 hours. Enterotoxigenic *Bacteroides fragilis* (ETBF) and non-enterotoxigenic *B. fragilis* (NTBF) were grown on TSA with 5% defibrinated sheep blood (TSBA) from skim milk stocks and incubated at 37°C anaerobically for 5 days. *Lactobacillus rhamnosus GG* was grown on TSA and incubated at 37°C for 24 hours aerobically and anaerobically.

For testing of various growth agar, *E. coli* DH5α was plated on TSA, TSBA, McConkey and eosin methylene blue agars. The plates were then incubated aerobically at 37°C for 24 hours. For testing of temperature stress environment, overnight cultures of DH5α in Lysogeny broth (LB) were diluted and incubated at 37°C or 50°C for 15 and 30 minutes. For testing of residual effects of environment, overnight cultures of DH5α in tryptic soy broth (TSB) were subcultured in various media and then plated on TSA. Briefly, DH5α overnight culture in TSB was centrifuged at 3,000 x g for 5 minutes and washed in sterile PBS twice. The pellet was then resuspended in PBS and cultured in LB at 37°C or 50°C for 1 hour, shaking. Samples were then plated on TSA plates and incubated at 37°C for 24 hours.

### Isolation and growth of clinical samples

Stool samples were collected from pre-diabetic adolescents aged 13-19 years as part of an ongoing study after institutional IRB approval. Participants were given a sterile stool collection kit (stool collection container [Precision (Covidien, Fisher Scientific)]), stool collector hat and sterile gloves to self-collect stool at home and mail to the University of Washington laboratory within 24-48 hours of collection at room temperature. When received, the sample was immediately aliquoted and one tube was used for the analysis of ‘Fresh’ sample on the same day it was received. The other aliquots were stored at −80°C for 2-3 months until analyzed. For analysis of the ‘Frozen sample’, the sample was thawed on ice for 30 mins and then room temperature for 30 mins. 40-80 mg of each sample suspended in 1ml PBS and then centrifuged at 300 rpm for 2 mins. Serial dilutions were made and plated on TSA and TSA blood and incubated in aerobic and anaerobic conditions at 37°C for 48 hours. Both ‘Fresh’ and ‘Frozen’ samples were analyzed using the direct protein method.

### Protein Extraction Method for MALDI-ToF MS

Bacterial colonies on agar were picked and added to 300 µl of HPLC-grade water in a 1.5 ml Eppendorf tube and then mixed thoroughly with 900 µl of 100% Ethanol. After centrifugation at 13000 rpm for 2 min twice, pellets were dried at room temperature for 5 minutes then they were directly mixed with equal volumes of 70% formic acid and acetonitrile (20-40 µl, depending on pellet size) for protein extraction. After centrifugation at 13000 rpm for 2 minutes, 1 µl of protein extract was spotted on a 96-target polished steel plate (Bruker Daltonics, Bremen, Germany) in four replicates, air-dried, and overlaid with 1 µl of alpha-cyano-4-hydroxycinnamic acid matrix solution (Bruker Daltonics, Bremen, Germany). Target plate was placed in the MALDI-ToF MS MicroFlex Biotyper for microbial identification (Bruker, Germany).

### Extended Direct Method for MALDI-ToF MS

Bacterial colonies on agar were picked and placed directly on a target plate for identification. Samples were then overlaid with 1 µL of 70% formic acid and alpha-cyano-4-hydroxycinnamic acid matrix solution (Bruker Daltonics, Bremen, Germany). Target plate was placed in the MALDI-ToF MicroFlex Biotyper for microbial identification (Bruker, Germany).

### Cellular adhesion and invasion assays with *Y. enterocolitica*

SW620s (ATCC; CCL-227) human intestinal epithelial adenoma cells were used. SW620s were maintained in DMEM supplemented with 10% fetal bovine serum and 200uM L-glutamine. Prior to cellular assays, SW620s were seeded into 24-well plates and grown to a nearly confluent monolayer over 3-4 days. SW620s were rinsed with PBS and infected with *Y. enterocolitica* at an MOI of 10:1 with each starting dose plated on TSA to enumerate CFUs. Briefly, overnight *Y. enterocolitica* cultures were diluted 1:10 in TSB. Subcultures were then incubated for 2 hours at 37°C while shaking at 150 rpm. Cultures were then centrifuged, rinsed in PBS and diluted to the appropriate infection dose in cell media. The *Y. enterocolitica* samples were introduced to the cells and infection initiated by centrifuging the plates for 5 minutes at 500 x g. For adhesion, Y*. enterocolitica* was co-cultured with SW620s for 1 hour at 37°C in a 5% CO_2_ incubator. For intracellular invasion, *Y. enterocolitica* was co-cultured with SW620s for 1 hour, washed with PBS and then treated with 100 µg/ml gentamicin (Corning, Corning, NY). To elute the *Y. enterocolitica* and lyse the SW620s, each well was washed three times with warm PBS and then treated with 500ul PBS with 1% Triton X-100 (AMRESCO, VWR, Randor PA) for 5 minutes. Samples were then plated on TSA in serial dilutions to enumerate *Y. enterocolitica* colonization. In parallel, wells containing media only were infected with an equal amount of bacteria. Percentage of colonization was calculated by dividing the number of *Y. enterocolitica* CFU recovered from co-cultures by the number of *Y. enterocolitica* CFU initially applied to the wells.

### Methods Data analysis

Raw spectra text files were analyzed using the R package, MALDIquant [https://www.ncbi.nlm.nih.gov/pubmed/22796955]. The raw data were trimmed to a spectra range of 3,000 to 15,000 m/z. The spectra intensities were then square-root transformed and smoothed using the Savitzky-Golay algorithm. Baseline noise was removed using the statistics-sensitive non-linear iterative peak clipping (or SNIP) algorithm with 100 iterations. The data were then normalized using total ion current (or TIC) calibration, which sets the total intensity to 1. Multiple spectra within the same analysis were aligned to the same x-axis using the Lowess warping method, a signal-to-noise ratio of 3, and a tolerance of 0.001. Peaks were detected from the average of at least 4 technical replicates using median absolute deviation. Principal components analyses and hierarchical clustering were also performed in R using the base stats package. Hierarchical clustering was performed on a calculated Euclidean distance matrix using Ward’s method.

## RESULTS AND DISCUSSION

### Peptide mass fingerprints distinguish genetic differences within microbial species

MALDI-ToF based bacterial profiling at the genus and species levels has provided results that are superior to and less expensive and time consuming than those obtained from more conventional approaches such as 16S rRNA sequencing. Such successes can be found across diverse areas of research and across many disciplines such as clinical microbiology, biodefense, food safety and environmental health. Although many studies have reported strain-specific peaks generated by MALDI-ToF MS, identification of reliable peaks as strain-specific biomarkers has been hindered by poor profile reproducibility. Further, it appears that the limits of the taxonomic resolution of MALDI-ToF MS profiling at the strain level has been determined so far in large part by the nature of the particular bacterium profiled. The more genetically indistinguishable bacteria are, the more challenging their profiling has been^28^. In this study, we show that MALDI-ToF MS offers the possibility to discriminate between highly genetically similar genetic mutants or between different strains of the same bacterial species. To investigate the capabilities of MALDI-ToF to distinguish strain-level differences, we analyzed three different gram-negative bacteria.

Two sample preparation methods have been mainly reported for MALDI-ToF MS-based identification and are known as the “Direct Transfer Method” and the “Protein extraction Method”^29^. The “Direct Transfer Method” consists in picking bacterial material from single colony forming units on a culture plate with a sterile transfer device and applying the material as a thin layer onto the MALDI-ToF target plate. The sample is then overlaid with a matrix for MALDI-ToF MS analysis^29^. While this method is simple and rapid, it is inferior in accuracy to the “Protein Extraction Method” because of insufficient cell wall disruption^29^. The “Protein Extraction Method” uses formic acid and acetonitrile to disrupt the cells prior to applying protein extracts directly to the target plate. In our experiments, it was proven that the “Protein Extraction Method” was necessary for the best performance in order to reliably and reproducibly detect strain level differences between the various tested microorganisms.

*Bacteroides fragilis* is a gram-negative, obligate anaerobic, rod-shaped bacterium. It is part of the normal microbiota of the human colon and is generally commensal but can cause infection if displaced into the bloodstream or surrounding tissue following surgery, disease, or trauma^30^. Enterotoxigenic *B. fragilis* (ETBF) strains are strains of *B. fragilis* that secrete a 20-kDa heat-labile zinc-dependent metalloprotease toxin termed the *B. fragilis* toxin (BFT). ETBF strains are associated with inflammatory diarrheal disease in children older than 1 year of age and in adults as well as in inflammatory bowel disease flare-ups and colorectal cancer^31–34^. In this experiment, the BFT positive strain of *B. fragilis* (BFT+) was grown alongside its mutant (BFT-) on TSA supplemented with 5% defibrinated sheep blood for 48 hours at 37°C anaerobically. Multiple colonies of the WT strain and mutant were tested in four technical replicates each on a MALDI-ToF target following protein extraction. A PCA was generated after analyzing the peptide mass fingerprint profiles of each colony, showing statistically significant differences between the proteomic profiles of the WT strain and mutant. The PMFs were generated using Bruker’s FlexAnalysis software and the PCA was performed with the help of the MALDIquant package in R. The mean PMF of the biological replicates from each cluster is shown by the larger point in the middle of each ellipse. This experiment clearly shows that the MALDI-ToF MS is sensitive enough to detect small genetic differences in highly similar microorganisms as it can reliably discriminate between the enterotoxigenic *B. fragilis* strain and a genetic mutant for the *B. fragilis* toxin (**Figure 1A**).

**Figure 1.**
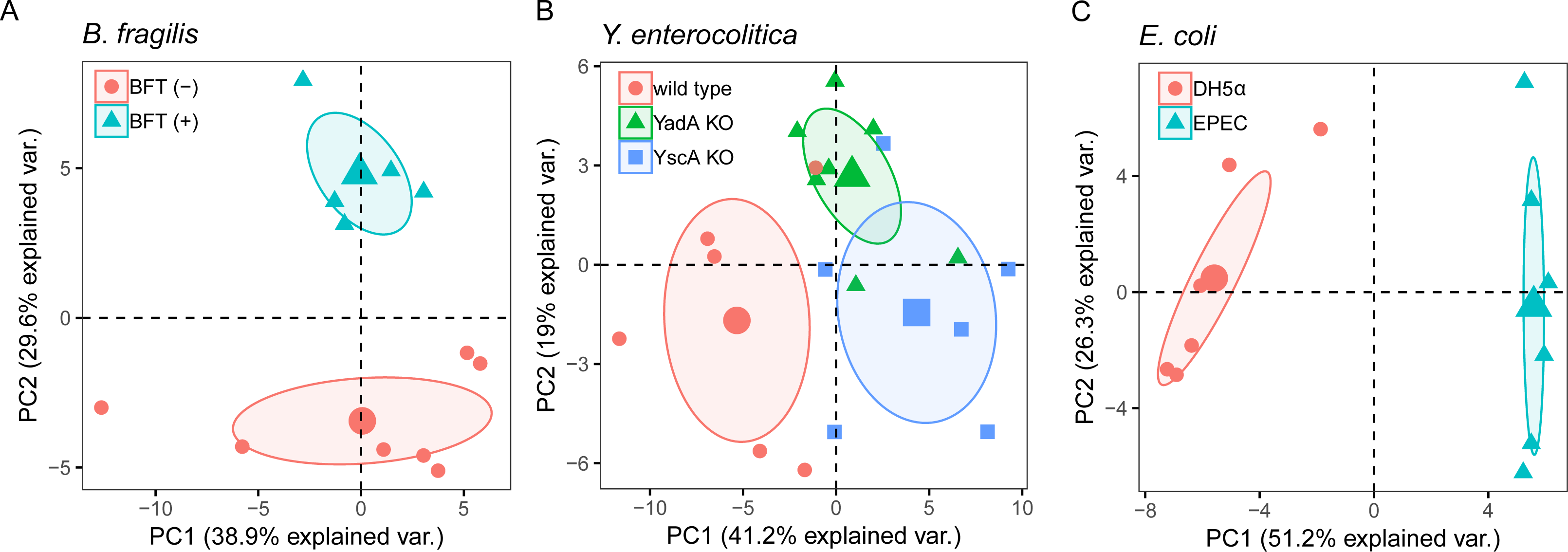
Differentiation of bacterial genetic mutants on MALDI-ToF MS. Principal component analysis (PCA) based on peptide mass fingerprint (PMF) profile of various bacterial strains and mutants. A) *Bacteroides fragilis* with and without the enterotoxigenic gene (BFT) are compared. B) *Yersinia enterocolitica* strain 8081 WT (Ye8081) and mutants with genetic differences in genes YadA and YscA are compared. C) Enteropathogenic *Eschericia coli* and DH5α strains are compared. Each dot represents the average of 4 technical replicates; the larger point is the mean of the experimental group with a 95% confidence interval drawn on mean. Data for each experiment are from two independent experiments with n=5-7.

Another gram-negative bacteria is *Yersinia enterocolitica. Y. enterocolitica* is a common food-borne disease found in contaminated water and meat. *Y. enterocolitica* is a transient infection in immunocompetent adults but can be deadly to immunocompromised individuals especially young children under the age of five^35, 36^. As before, the various strains of *Y. enterocolitica* were grown on simultaneously prior to protein extraction and analysis through MALDI-ToF MS. The three strains of *Y. enterocolitica* were grown on TSA for 48 hours aerobically at room temperature. Using the MALDI-ToF, we are able to see three distinct clusters among a wild type *Y. enterocolitica* strain (Ye8081) and two mutants of the same strain that are lacking an adhesion gene, *YadA*, or a *Yersinia* translocation protein *(YscA)* (**Figure 1B**).

*Escherichia coli* is a gram-negative, rod-shaped bacterium that is commonly found in the lower intestines. While most *E. coli* strains are harmless, pathogenic varieties can cause serious gastroenteritis, urinary tract infections, meningitis or septic shock in humans^37^. Enteropathogenic *E. coli* (EPEC) is a strain of *E. coli* that uses a virulence factor known as intimin, an adhesin that binds host intestinal cells, causing watery diarrhea in those afflicted^38^. In this experiment, EPEC was grown alongside a non-pathogenic strain of *E. coli*, DH5α on TSA for 24 hours at 37°C aerobically. As we demonstrated with the *B. fragilis* and *Y. enterocolitica*, the MALDI-ToF can also differentiate lab strains of *E. coli* from the highly pathogenic EPEC (**Figure 1C**). This experiment clearly shows that MALDI-ToF MS is sensitive enough to detect strain-level differences within the same bacterial species as it was able to differentiate lab strains of *E. coli* from the highly pathogenic EPEC.

As the MALDI-ToF MS uses whole protein of bacterial samples to generate unique profiles, lack of phenotypic or metabolic differences between subgroups of species can complicate identification. Meanwhile, our results confirm the findings of other studies. The technique has proved to be highly performant and reproducible in distinguishing peaks within highly genetically similar species. Several known strains can then be used to create a reference library that affords identification of unidentified strains with potentially useful applications such as diagnosing pathogenic strains in certain disease states or rapidly identifying the origin of an outbreak.

### PMFs distinguish phenotypic differences within genetically identical strains

The reproducibility of the mass spectra generated in MALDI-ToF MS analysis of proteins from bacterial extracts can be impacted by several experimental factors. Other researchers have discovered that differences in incubation and culturing conditions can alter peak intensities^39^ and identification rates^40^ (**Table 1**). In order to test whether environmental differences could reproducibly impact the proteome of several microorganisms, we decided to look at various environmental factors such as aerobic versus anaerobic conditions, various culturing medias, and different incubation temperatures on how they impact the PMF profile of the tested microorganisms.

**Table 1.**
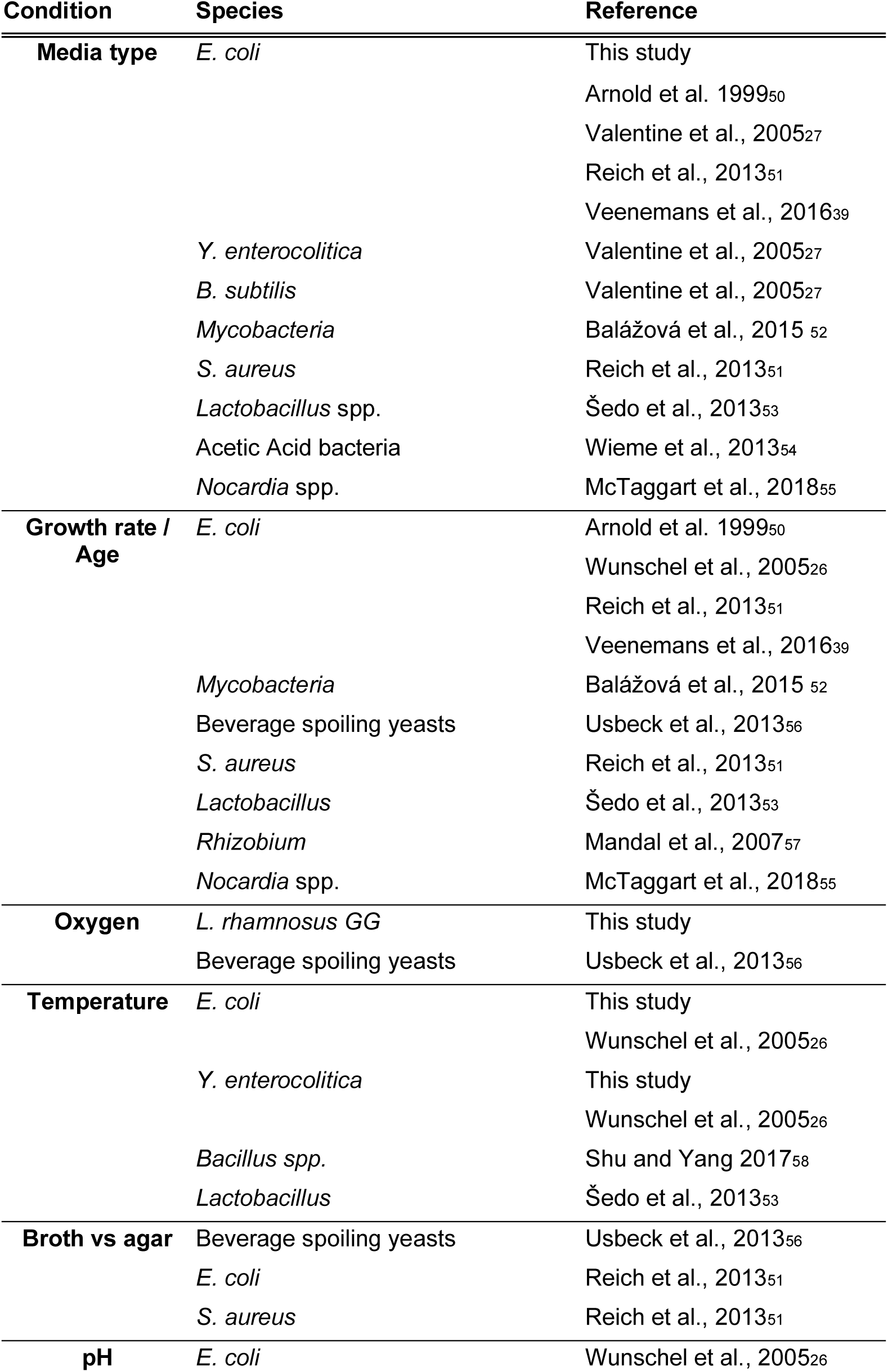
Environmental influences on MALDI-ToF MS analyses

The first environmental factor tested was presence or absence of oxygen. Using the microorganism *Lactobacillus reuteri*, we investigated if MALDI-ToF MS analysis could detect differences in PMF following overnight incubation at 37°C in aerobic and anaerobic environments. *Lactobacillus spp.* are facultative anaerobes and several genes are differently expressed with the presence of oxygen^41^. Consequently, our experiment reveals that despite using the same starting culture, aerobic or anaerobic conditions cause significant alterations in the PMF profile of the same microbial strain (**Figure 2A**). The two conditions clearly separate along the x-axis with over 50% of the various explained by PC1 (**Figure 2A**). Interestingly, the clustering of *L reuteri* grown aerobically exhibit more variability in PMFs compared to *L. reuteri* grown anaerobically. These differences may be indicative of the impact of anaerobic stress exerting a more distinct response on the bacteria.

**Figure 2.**
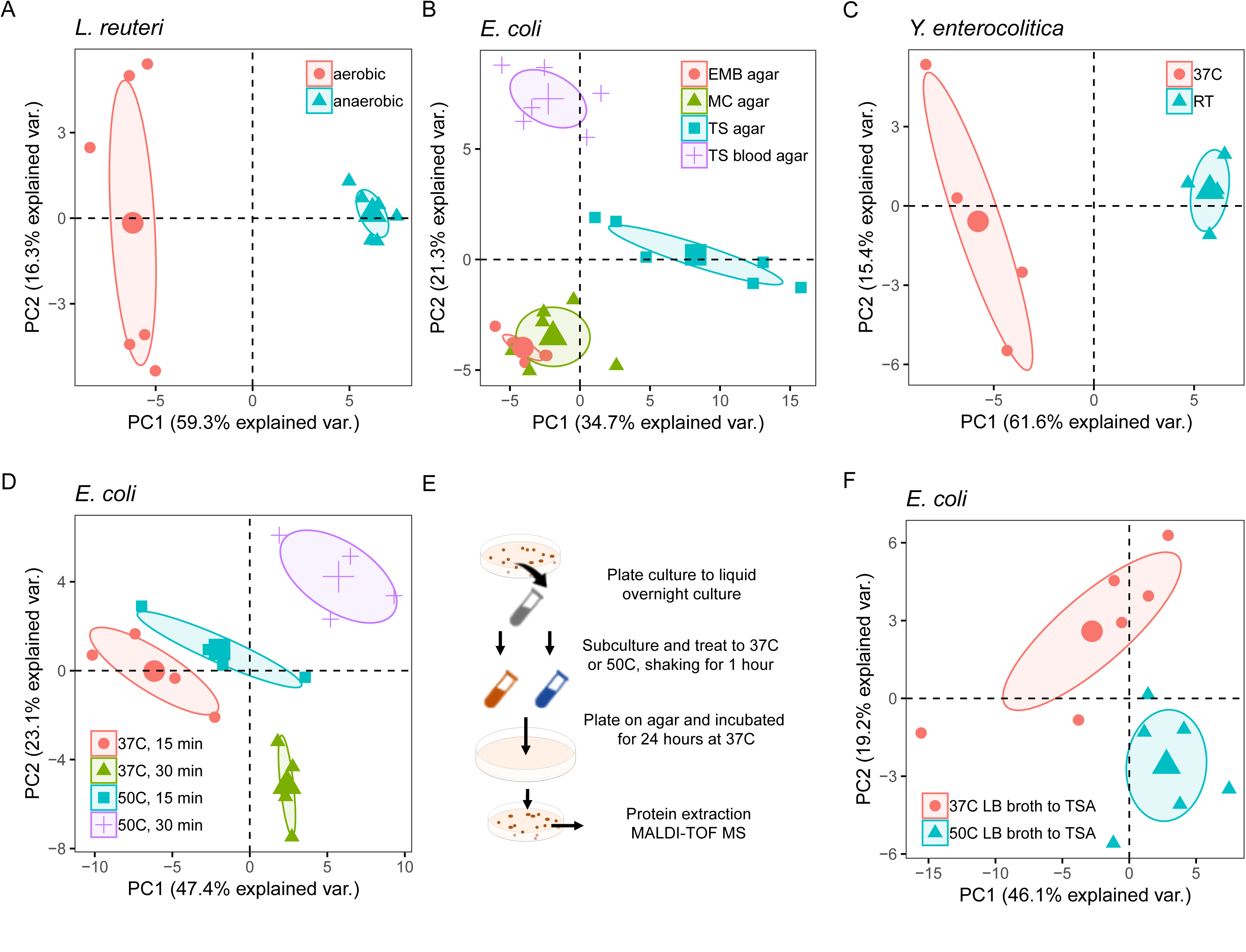
Differentiation of bacterial biological environment on MALDI-ToF MS. PCA based on PMF profile of various bacterial species cultured in different growth conditions. A) *Lactobacillus reuteri* plated on TSA and incubated overnight at 37°C in aerobic and anaerobic conditions is compared. B) DH5α *Escherichia coli* strain grown on 4 different types of Media: TSA (TS agar), TSA with 5% sheep blood (TS blood agar), McConkey agar (MC agar), and eosin methylene blue agar (EMB agar) is compared. C) *Y. enterocolitica* grown at room temperature (RT) or 37°C for 48 hours on TSA. D) DH5α grown in liquid culture at 37°C and 50°C for 15 and 30 min while shaking. E) Diagram and F) PCA of DH5α grown in liquid culture at 37°C and 50°C for 1 hour and then plated to observe residual effects of different environmental conditions. Each dot represents the average of 4 technical replicates; the larger point is the mean of the experimental group with a 95% confidence interval drawn on mean. Data for each experiment are from two independent experiments with n=5-7.

As others have noted, culturing on various media does not alter species identification. For this study we cultured *E. coli* DH5α on four different nutrient-rich media: TSA, TSA supplemented with 5% blood (TSBA), MacConkey agar (MC), and eosin methylene blue agar (EMB). Whereas *E. coli* grown on MC and EMB agar had overlapping PMF profiles and clustered together on the PCA, there were clearly different clusters for *E. coli* grown on TSA and TSBA. (**Figure 2B**). The overlapping of MC and EMB-grown *E. coli* is likely due to the lactose fermentation process undergone by *E. coli* when it is grown on these two types of agars. While the differences between MC and EMB agars did not discriminate between the *E. coli* colonies grown on these two types of media, the presence of 5% defibrinated sheep blood in the TSA was enough to trigger the expression of a different proteomic profile in *E. coli* grown on TSBA relative to TSA (**Figure 2B**). Thus the MALDI-ToF MS has the capacity to significantly discriminate between *E. coli* grown on various media depending on the components of the media.

We next tested whether temperature affects the PMF profiles of microorganisms. Gut pathogen *Y. enterocolitica* is well adapted for survival and proliferation at room temperature (RT) and induces its virulence factors when consumed or introduced to host temperatures of 37°C^42^. These metabolic adaptations are clearly captured when we analyze *Y. enterocolitica* grown at these two temperatures on the MALDI-ToF MS with *Y. enterocolitica* grown at RT clustering away from the 37°C treated group (**Figure 2C**). The two clusters for *Y. enterocolitica* are clearly separated on the x-axis with over 60% of the PMF various explained on PC1, demonstrating a clear distinction in proteome expressions due to temperature conditions. The individual PMFs of *Y. enterocolitica* grown at room temperature are also more tightly clustered than the *Y. enterocolitica* grown at 37°C. These clustering could be indicative of more variable protein expression of *Y. enterocolitica* when grown in virulence-inducing temperatures. Previous studies using MALDI-ToF MS on *Y. enterocolitica* point out that temperature and media both play a role in PMF intensities with inference that temperature may play a bigger role^26, 27^. Whereas Wunschel and collegues^26^ identified that *Y. enterocolitica* grown at 37°C has similar PMFs despite the media, we demonstrate that the MALDI-ToF can reliably distinguish between the temperature conditions.

Next we tested whether stressing a bacteria with high heat induces heat shock proteins^43^. For this experiment, liquid cultures of DH5α were incubated at 37°C or 50°C for 15 minutes or 30 minutes. Sure enough, we clearly saw that temperatures also affected the PMF profile and clustering of *E. coli* introduced to temperatures greater than 42°C. These differences, most likely due to the translation of heat-shock proteins, became even more dramatic with an increase in incubation time (**Figure 2D**).

All together these results with bacteria grown in different oxygen, nutrient and temperature environments demonstrate how the MALDI-ToF is capable of reproducibly discriminating between metabolic states through PMF profiles. Other studies have investigated whether these environmental and culture conditions can impact the MALDI-ToF identification of bacteria. Our experiments took it one step further and demonstrate that the MALDI-ToF can easily differentiate the direct effects of environmental conditions on the bacteria, allowing identification of biological environmental influences as it affects the proteomic profile of microorganisms.

To further understand the extent of the MALDI-ToF MS technique on discerning environmental influences on bacteria, we tested if previous environmental stressors can impact PMF clustering. For this experiment, DH5α was grown in liquid culture at 37°C and 50°C for 2 hour and then plated on TSA for 24 hours. Whereas our previous experiment measured the direct environmental impact of temperature on DH5α (**Figure 2D**), this experiment examined if the MALDI-ToF could sense the same temperature effects after the bacteria was removed from that conditions and plated. As expected, the PMF profiles of DH5α grown in the two different cultures had distinct clusters (**Figure 2E-F**). While there were some overlaps in clusters, this data demonstrates that the MALDI-ToF can identify environmental conditioning effects despite the removal of the immediate conditions. These results have major implications in the lab setting where microorganisms isolated from biological samples are often studied *ex vivo*. Our ability to detect environmental conditionings even when the bacteria is removed from the initial stresses further highlight the advantage of proteotyping using the MALDI-ToF MS.

### PMFs identify metabolic and virulent states of *Y. enterocolitica* during in vitro adhesion and invasion assay

MALDI-TOF analysis of *Y. enterocolitica* at the various temperatures demonstrated that PMF can potentially distinguish between virulent states. Other than temperature, *Y. enterocolitica* has different metabolic states during infection and various genes that are modified during the course of infection^44^. The stages of *Y. enterocolitica* infection were investigated using human intestinal cells, SW620s. PMFs of untreated *Y. enterocolitica*, *Y. enterocolitica* applied to SW620s for 1 hour, or applied to SW620s for 1 hour and treated with gentamicin to remove extracellular bacteria were compared (**Figure 3A**). After samples were collected, *Y. enterocolitica* was plated on TSA and incubated at room temperature for 48 hours.

**Figure 3.**
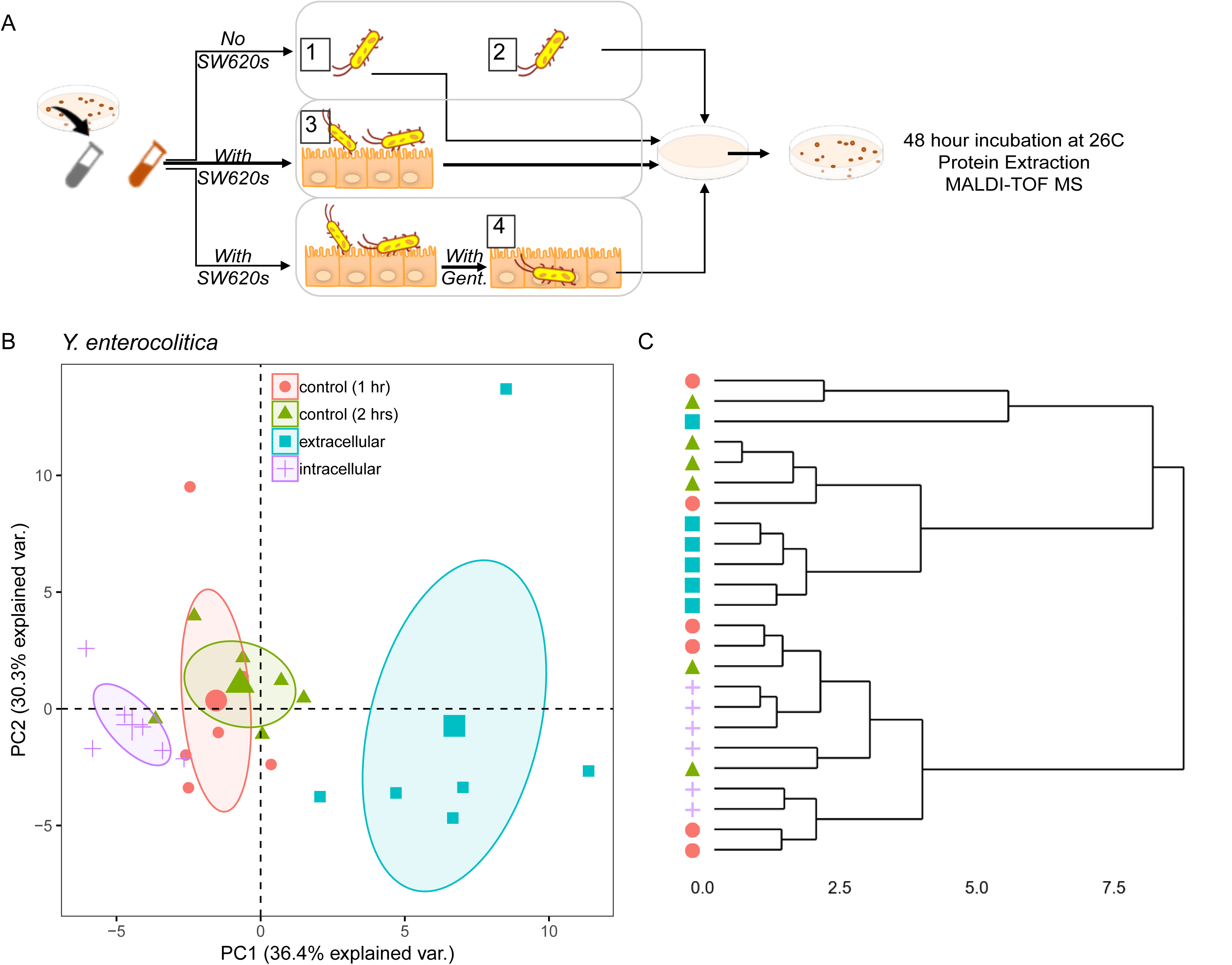
PMF analysis of *Y. enterocolitica* virulence states. A) Schematic of protocol. B) PCA and dendrogram of *Y. enterocolitica* cultured 48 hours at room temperature on TSA following adhesion and invasion assay with SW620 cells. Each dot represents the average of 4 technical replicates; the larger point is the mean of the experimental group with a 95% confidence interval drawn on mean. Data for each experiment are from two independent experiments with n=6.

As we previously demonstrated with liquid cultures of DH5α plated on TSA, we found that *Y. enterocolitica* clustered depending on treatment group (**Figure 3B**). Cultures of intracellular *Y. enterocolitica* recovered following gentamicin had greater similarity whereas the *Y. enterocolitica* recovered after 1 hour had a more variable PMF profile. More divergent PMFs within a population could be due to natural heterogeneity within a clonal population^45^ or differences in transcriptional or virulent states.

Our data using DH5α *E. coli* and *Y. enterocolitica* demonstrate that the MALDI-ToF can detect differences in metabolic and virulence states, respectively, even after those environmental influences are removed. This is especially important for trying to assess phenotypic differences of clinical isolates or samples that cannot be readily grown in its natural environments. These methods could be applied to situations in which metabolic or phenotypic difference of experimental groups can be compared against known culturing conditions. It can be used as a method of screening of known bacterial phenotypes.

### PMFs distinguish between fresh and frozen clinical isolates of the same species

A very common use of the MALDI-ToF MS in clinical studies is to identify clinical isolates removed from biological samples for diagnostic purposes. Whereas the previous experiments described above used the protein extraction method in a well-controlled environment with well characterized strains, the use of MALDI-ToF in clinical settings is less likely to resort to this time-consuming method. In a clinical setting, using the protein extraction method is time consuming and not all samples are easily processed immediately. Additionally, freezing samples prior to culturing can reduce the number of bacteria recovered, skew the diversity of bacteria, and alter community composition^46, 47^. Therefore these next experiments evaluated how frozen versus fresh conditions altered the PMFs of clinical samples using the “Extended Direct Transfer” method as described above.

Using fresh human fecal samples, we tested whether culturing fresh or frozen microbiota samples altered the phenotype of clinical isolates (**Figure 4A**). In these experiments, whole fecal samples were plated aerobically on TSBA for 48 hours. Single colonies were then processed using the extended direct colony method. Bacteria identified as the same species were then compiled and examined retrospectively.

**Figure 4.**
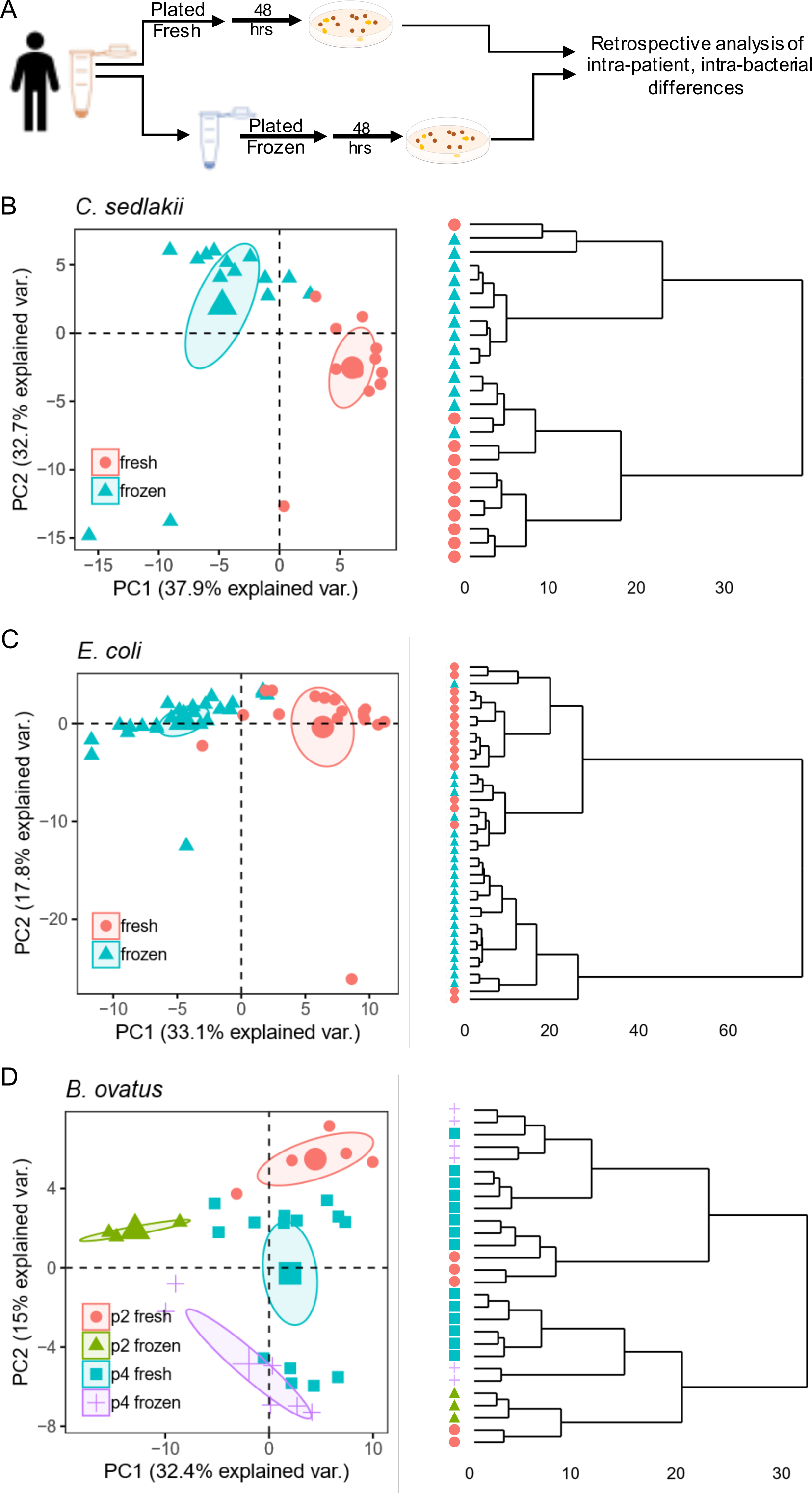
PMF analysis of fresh and frozen clinical isolates. A) Schematic of clinical sample analysis. PCA and dendrogram of fresh and frozen PMFs from B) *C. sedlakii*, C) *E. coli* and D) *B. ovatus* isolates identified on aerobic TSA with 5% sheep blood. Isolates proceeded via extended direct transfer. Each dot resembles a biological replicate. Ellipse drawn on the 95% confidence interval of the mean.

Within our clinical samples, the MALDI-ToF identified *Citrobacter sedlakii*, *E. coli* and *Bacteroides ovatus* from both fresh and frozen cultures. Despite distinct clustering between fresh and frozen isolates of *C. sedlakii* and *E. coli*, the clustering was not statistically significant as there was overlap seen in the PCA and dendrogram (**Figure 4B-C**). Freezing seemed to have a more profound impact on *C. sedlakii* compared to *E. coli* as these isolates appeared to have little overlap in PMFs (**Figure 4B**). In contrast, *E. coli* isolates from fresh and frozen stool had more similarities suggesting that the response to freezing may be bacteria dependent (**Figure 4C**). These experiments provide a rationale for why sample collection during microbiome studies must be standardized.

We next compared the MALDI-ToF analysis of fresh and frozen *B. ovatus* taken from two different patients clustered. As expected, *B. ovatus* clustering was patient dependent, likely due to each patient containing a different strain of *B. ovatus* or exposure to patient dependent microenvironments that contribute to genetic diversity via phage and horizontal gene transfer (**Figure 4D**). Interestingly, fresh or frozen status had less of an impact on the clustering of *B. ovatus* from patient 2, while patient 1 had high variability of PMFs within fresh and frozen groups. The different sensitivity between patient 1 and patient 2 *B. ovatus* isolates from fresh and frozen stool may be due to the presence of more temperature sensitive genes in the isolates from patient 1 and indicate that freezing stool may alter the PMF of MALDI-ToF in a bacteria-dependent manner.

Our results using clinical isolates derived from a mixed microbiota sample reveal that fresh versus frozen culturing techniques may alter the proteotyping of microorganisms. Further study is needed to analyze the freezing effects. This clinical experiment was limited by MALDI-ToF processing, lack of technical replicates, and difference in MALDI-ToF sampling days. These results exemplify a need for rigorous sample preparation and analysis. Given our ability to proteotype bacteria by various environmental factors, there is huge potential in analyzing phenotypic differences amongst clinical samples. There are numerous applications should this methodology be adapted in the clinic such as distinguishing proteotypes from inflammatory, nutrient-rich or -depleted, or competitive microenvironments.

## CONCLUSIONS

The emergence of MALDI-ToF MS has reinvigorated microbial identification over the past decade. By creating proteomic signatures through PMFs of microorganisms, species and subspecies identification is possible via proteotyping. Routine use of MALDI-ToF MS as a diagnostic tool is favorable due to its ease of use, time and cost efficiency, and robust sensitivity. Our current study demonstrates a new function by exploiting the unique proteomic signatures and comparative analysis afforded by the MALDI-ToF MS bioinformatic packages. While proteotyping of different physiological and metabolic states of microorganisms through MALDI-ToF has been previously suggested by others^48, 49^ and variations in culturing conditions have been extensively tested for impact on species identification (Table 1), we test the ability to distinguish between multiple environmental conditions within the same microorganism. Environmental effects of oxygen, temperature and nutrients can all be easily distinguished. As a proof-of-principle, we also demonstrate that PMF signatures can differentiate by stressors even when microorganisms are removed from the initial environment. We further exemplify how the MALDI-ToF MS technique can separate different virulent states of *Y. enterocolitica* with the SW620 adhesion assay.

One limitation of proteotyping of metabolic and functional states include lack of PMF peak to protein identification readily available through the Bruker software. While publicly accessible databases for peptide/protein identification and assignment exist, such as Mascot (www.matrixscience.com), there still remains a need to develop a database of peak information that can be more readily incorporated with MALDI-ToF MS data. Another limitation is addressed with our clinical data in which rigorous protein extraction and technical replicates may not be easily incorporated in a clinical setting. While our data inconclusively delineate fresh versus frozen isolates, these experiments need to be repeated with a wide variety of microorganisms.

Altogether, we demonstrate the potential of MALDI-ToF MS to screen for physiological changes from direct and indirect environmental conditions. In conjunction with transcriptomic data and validation through PCR or LC/MS, this technique could provide a rapid and efficient way to screen for metabolic functions and physiological states of microorganisms. This methodology can be especially useful for analyzing phenotypic changes of bacteria cultured from previously unique environments such as an inflamed gastrointestinal tract or progression through virulence. In addition to the original function of species identification, the MALDI-ToF MS provides a plethora of data that has yet to be fully explored and exploited. Pairing these data with peptide/protein identity assignment could ultimately constitute a strong framework for the identification of microbiome-based biomarkers and therapeutic targets in various disease states.

## ACKNOWLEDGMENTS

The authors thank Dr. Marion Avril for support and proofreading. The authors thank the Center for Microbiome Sciences & Therapeutics (CMiST) at the University of Washington for the MALDI-ToF MS resources. The authors thank Dr. Cynthia Sears for graciously providing the wild type and BFT-negative *B. fragilis* strains.

## REFERENCES

1. Singhal, N., Kumar, M., Kanaujia, P. K. & Virdi, J. S. MALDI-TOF mass spectrometry: an emerging technology for microbial identification and diagnosis. Front. Microbiol. 6, (2015).

2. Bhavsar, S. M., Dingle, T. C. & Hamula, C. L. The impact of blood culture identification by MALDI-TOF MS on the antimicrobial management of pediatric patients. Diagnostic Microbiology and Infectious Disease 92, 220–225 (2018).

3. Khan, S. et al. MALDI-TOF MS in Adult Inpatients with Bloodstream Infections: Pre- and Post-intervention Study. Open Forum Infectious Diseases 4, S626–S626 (2017).

4. Haiko, J., Savolainen, L. E., Hilla, R. & Pätäri-Sampo, A. Identification of urinary tract pathogens after 3-hours urine culture by MALDI-TOF mass spectrometry. Journal of Microbiological Methods 129, 81–84 (2016).

5. Ferreira, L. et al. Direct Identification of Urinary Tract Pathogens from Urine Samples by Matrix-Assisted Laser Desorption Ionization-Time of Flight Mass Spectrometry. Journal of Clinical Microbiology 48, 2110–2115 (2010).

6. Íñigo, M. et al. Direct Identification of Urinary Tract Pathogens from Urine Samples, Combining Urine Screening Methods and Matrix-Assisted Laser Desorption Ionization-Time of Flight Mass Spectrometry. J. Clin. Microbiol. 54, 988–993 (2016).

7. Svarrer, C. W. & Uldum, S. A. The occurrence of Legionella species other than Legionella pneumophila in clinical and environmental samples in Denmark identified by mip gene sequencing and matrix-assisted laser desorption ionization time-of-flight mass spectrometry. Clinical Microbiology and Infection 18, 1004–1009 (2012).

8. Samb-Ba, B. et al. MALDI-TOF Identification of the Human Gut Microbiome in People with and without Diarrhea in Senegal. PLoS ONE 9, e87419 (2014).

9. Rizzardi, K. & Åkerlund, T. High Molecular Weight Typing with MALDI-TOF MS -A Novel Method for Rapid Typing of Clostridium difficile. PLoS ONE 10, e0122457 (2015).

10. Wang, H.-Y. et al. A new scheme for strain typing of methicillin-resistant Staphylococcus aureus on the basis of matrix-assisted laser desorption ionization time-of-flight mass spectrometry by using machine learning approach. PLoS ONE 13, e0194289 (2018).

11. Wolters, M. et al. MALDI-TOF MS fingerprinting allows for discrimination of major methicillin-resistant Staphylococcus aureus lineages. International Journal of Medical Microbiology 301, 64–68 (2011).

12. Manukumar, H. M. & Umesha, S. MALDI-TOF-MS based identification and molecular characterization of food associated methicillin-resistant Staphylococcus aureus. Sci Rep 7, 11414 (2017).

13. Nisa, S. et al. Combining MALDI-TOF and genomics in the study of methicillin resistant and multidrug resistant Staphylococcus pseudintermedius in New Zealand. Sci Rep 9, 1271 (2019).

14. Christner, M. et al. Rapid MALDI-TOF Mass Spectrometry Strain Typing during a Large Outbreak of Shiga-Toxigenic Escherichia coli. PLoS ONE 9, e101924 (2014).

15. Alanio, A. et al. Matrix-assisted laser desorption ionization time-of-flight mass spectrometry for fast and accurate identification of clinically relevant Aspergillus species. Clinical Microbiology and Infection 17, 750–755 (2011).

16. Tenover, F. C., Arbeit, R. D. & Goering, R. V. How to select and interpret molecular strain typing methods for epidemiological studies of bacterial infections: a review for healthcare epidemiologists. Molecular Typing Working Group of the Society for Healthcare Epidemiology of America. Infect Control Hosp Epidemiol 18, 426–439 (1997).

17. Parizad, E. G., Parizad, E. G. & Valizadeh, A. The Application of Pulsed Field Gel Electrophoresis in Clinical Studies. J Clin Diagn Res 10, DE01–DE04 (2016).

18. Killgore, G. et al. Comparison of seven techniques for typing international epidemic strains of Clostridium difficile: restriction endonuclease analysis, pulsed-field gel electrophoresis, PCR-ribotyping, multilocus sequence typing, multilocus variable-number tandem-repeat analysis, amplified fragment length polymorphism, and surface layer protein A gene sequence typing. J. Clin. Microbiol. 46, 431–437 (2008).

19. de Bruijn, F. J. Use of repetitive (repetitive extragenic palindromic and enterobacterial repetitive intergeneric consensus) sequences and the polymerase chain reaction to fingerprint the genomes of Rhizobium meliloti isolates and other soil bacteria. Appl. Environ. Microbiol. 58, 2180–2187 (1992).

20. Parolo, C. C. F. et al. Genetic diversity of Lactobacillus paracasei isolated from in situ human oral biofilms. J. Appl. Microbiol. 111, 105–113 (2011).

21. Balloux, F. et al. From Theory to Practice: Translating Whole-Genome Sequencing (WGS) into the Clinic. Trends Microbiol 26, 1035–1048 (2018).

22. Schürch, A. C., Arredondo-Alonso, S., Willems, R. J. L. & Goering, R. V. Whole genome sequencing options for bacterial strain typing and epidemiologic analysis based on single nucleotide polymorphism versus gene-by-gene-based approaches. Clinical Microbiology and Infection 24, 350–354 (2018).

23. Murray, P. R. Matrix-assisted laser desorption ionization time-of-flight mass spectrometry: usefulness for taxonomy and epidemiology. Clin. Microbiol. Infect. 16, 1626–1630 (2010).

24. Elinav, E., Garrett, W. S., Trinchieri, G. & Wargo, J. The cancer microbiome. Nat Rev Cancer 19, 371–376 (2019).

25. Lasch, P. et al. Insufficient discriminatory power of MALDI-TOF mass spectrometry for typing of Enterococcus faecium and Staphylococcus aureus isolates. Journal of Microbiological Methods 100, 58–69 (2014).

26. Wunschel, D. S. et al. Effects of varied pH, growth rate and temperature using controlled fermentation and batch culture on Matrix Assisted Laser Desorption/Ionization whole cell protein fingerprints. Journal of Microbiological Methods 62, 259–271 (2005).

27. Valentine, N., Wunschel, S., Wunschel, D., Petersen, C. & Wahl, K. Effect of Culture Conditions on Microorganism Identification by Matrix-Assisted Laser Desorption Ionization Mass Spectrometry. Applied and Environmental Microbiology 71, 58–64 (2005).

28. Ghyselinck, J., Van Hoorde, K., Hoste, B., Heylen, K. & De Vos, P. Evaluation of MALDI-TOF MS as a tool for high-throughput dereplication. Journal of Microbiological Methods 86, 327–336 (2011).

29. Alatoom, A. A., Cunningham, S. A., Ihde, S. M., Mandrekar, J. & Patel, R. Comparison of Direct Colony Method versus Extraction Method for Identification of Gram-Positive Cocci by Use of Bruker Biotyper Matrix-Assisted Laser Desorption Ionization-Time of Flight Mass Spectrometry. Journal of Clinical Microbiology 49, 2868–2873 (2011).

30. Wexler, H. M. Bacteroides: the Good, the Bad, and the Nitty-Gritty. Clinical Microbiology Reviews 20, 593–621 (2007).

31. Rashidan, M. et al. Detection of B. fragilis group and diversity of bft enterotoxin and antibiotic resistance markers cepA, cfiA and nim among intestinal Bacteroides fragilis strains in patients with inflammatory bowel disease. Anaerobe 50, 93–100 (2018).

32. Zamani, S. et al. Detection of enterotoxigenic Bacteroides fragilis in patients with ulcerative colitis. Gut Pathog 9, 53 (2017).

33. Purcell, R. V. et al. Colonization with enterotoxigenic Bacteroides fragilis is associated with early-stage colorectal neoplasia. PLoS ONE 12, e0171602 (2017).

34. Goodwin, A. C. et al. Polyamine catabolism contributes to enterotoxigenic Bacteroides fragilis-induced colon tumorigenesis. Proceedings of the National Academy of Sciences 108, 15354–15359 (2011).

35. Lee, L. A. et al. Yersinia Enterocolitica O:3: An Emerging Cause of Pediatric Gastroenteritis in the United States. Journal of Infectious Diseases 163, 660–663 (1991).

36. Ray, S. M. et al. Population-Based Surveillance for *Yersinia enterocolitica* Infections in FoodNet Sites, 1996-1999: Higher Risk of Disease in Infants and Minority Populations. CLIN INFECT DIS 38, S181–S189 (2004).

37. Chart, H., Smith, H. R., La Ragione, R. M. & Woodward, M. J. An investigation into the pathogenic properties of Escherichia coli strains BLR, BL21, DH5αlpha and EQ1. J. Appl. Microbiol. 89, 1048–1058 (2000).

38. Clarke, S. C., Haigh, R. D., Freestone, P. P. E. & Williams, P. H. Virulence of Enteropathogenic Escherichia coli, a Global Pathogen. Clinical Microbiology Reviews 16, 365–378 (2003).

39. Veenemans, J. et al. Comparison of MALDI-TOF MS and AFLP for strain typing of ESBL-producing Escherichia coli. Eur J Clin Microbiol Infect Dis 35, 829–838 (2016).

40. Vithanage, N. R. et al. Species-Level Discrimination of Psychrotrophic Pathogenic and Spoilage Gram-Negative Raw Milk Isolates Using a Combined MALDI-TOF MS Proteomics-Bioinformatics-based Approach. J. Proteome Res. 16, 2188–2203 (2017).

41. Zotta, T., Parente, E. & Ricciardi, A. Aerobic metabolism in the genus *Lactobacillus* : impact on stress response and potential applications in the food industry. J Appl Microbiol 122, 857–869 (2017).

42. Straley, S. C. & Perry, R. D. Environmental modulation of gene expression and pathogenesis in Yersinia. Trends Microbiol. 3, 310–317 (1995).

43. Tomoyasu, T. et al. Escherichia coli FtsH is a membrane-bound, ATP-dependent protease which degrades the heat-shock transcription factor sigma 32. EMBO J. 14, 2551–2560 (1995).

44. Bent, Z. W. et al. Transcriptomic Analysis of Yersinia enterocolitica Biovar 1B Infecting Murine Macrophages Reveals New Mechanisms of Extracellular and Intracellular Survival. Infect. Immun. 83, 2672–2685 (2015).

45. Ackermann, M. A functional perspective on phenotypic heterogeneity in microorganisms. Nat Rev Microbiol 13, 497–508 (2015).

46. Lau, J. T. et al. Capturing the diversity of the human gut microbiota through culture-enriched molecular profiling. Genome Med 8, 72 (2016).

47. Wu, W.-K. et al. Optimization of fecal sample processing for microbiome study - The journey from bathroom to bench. Journal of the Formosan Medical Association 118, 545–555 (2019).

48. Grenga, L., Pible, O. & Armengaud, J. Pathogen proteotyping: A rapidly developing application of mass spectrometry to address clinical concerns. Clinical Mass Spectrometry S2376999818300503 (2019) doi:10.1016/j.clinms.2019.04.004.

49. Karlsson, R. et al. Proteotyping: Proteomic characterization, classification and identification of microorganisms - A prospectus. Systematic and Applied Microbiology 38, 246–257 (2015).

50. Arnold, R. J., Karty, J. A., Ellington, A. D. & Reilly, J. P. Monitoring the Growth of a Bacteria Culture by MALDI-MS of Whole Cells. Anal. Chem. 71, 1990–1996 (1999).

51. Reich, M. Species Identification of Bacteria and Fungi from Solid and Liquid Culture Media by MALDI-TOF Mass Spectrometry. J Bacteriol Parasitol 01, (2013).

52. Balazova, T. et al. The influence of culture conditions on the identification of *Mycobacterium* species by MALDI-TOF MS profiling. FEMS Microbiol Lett 353, 77–84 (2014).

53. Sedo, O., Vavrova, A., Vad’urova, M., Tvrzova, L. & Zdráhal, Z. The influence of growth conditions on strain differentiation within the *Lactobacillus acidophilus* group using matrix-assisted laser desorption/ionization time-of-flight mass spectrometry profiling: Growth conditions influence strain differentiation using MALDI-TOF MS. Rapid Commun. Mass Spectrom. 27, 2729–2736 (2013).

54. Wieme, A. D. et al. Effects of Growth Medium on Matrix-Assisted Laser Desorption-Ionization Time of Flight Mass Spectra: a Case Study of Acetic Acid Bacteria. Appl. Environ. Microbiol. 80, 1528–1538 (2014).

55. McTaggart, L. R., Chen, Y., Poopalarajah, R. & Kus, J. V. Incubation time and culture media impact success of identification of Nocardia spp. by MALDI-ToF mass spectrometry. Diagnostic Microbiology and Infectious Disease 92, 270–274 (2018).

56. Usbeck, J. C., Kern, C. C., Vogel, R. F. & Behr, J. Optimization of experimental and modelling parameters for the differentiation of beverage spoiling yeasts by Matrix-Assisted-Laser-Desorption/Ionization-Time-of-Flight Mass Spectrometry (MALDI-TOF MS) in response to varying growth conditions. Food Microbiology 36, 379–387 (2013).

57. Mandal, S. M., Pati, B. R., Ghosh, A. K. & Das, A. K. Letter: Influence of Experimental Parameters on Identification of Whole Cell *Rhizobium* by Matrix-Assisted Laser Desorption/Ionization Time-of-Flight Mass Spectrometry. Eur J Mass Spectrom (Chichester) 13, 165–171 (2007).

58. Shu, L.-J. & Yang, Y.-L. Bacillus Classification Based on Matrix-Assisted Laser Desorption Ionization Time-of-Flight Mass Spectrometry-Effects of Culture Conditions. Sci Rep 7, 15546 (2017).

